# Optical opening of the blood brain barrier for targeted and ultra-sparse viral infection of cells in the mouse cortex

**DOI:** 10.1101/2022.11.10.516054

**Authors:** Patrick L. Reeson, Roobina Boghozian, Ana-Paula Cote, Craig E. Brown

## Abstract

Adeno-associated viruses (AAVs) are used in a wide array of experimental situations for driving expression of biosensors, recombinases and opto/chemo-genetic actuators in the brain. However, conventional approaches for minimally invasive, spatially precise and ultra-sparse AAV mediated transduction of cells during imaging experiments, has remained a significant challenge. Here we show that intravenous injection of commercially available AAVs at different doses, combined with laser based perforation of single cortical capillaries through a cranial widow, allows for ultra-sparse, titrate-able, and micron level precision for delivery of viral vectors with relatively little inflammation or tissue damage. Further, we show the utility of this approach for eliciting sparse expression of GCaMP6, channel-rhodopsin or fluorescent reporters in neurons and astrocytes within specific functional domains in normal and stroke damaged cortex. This technique represents a facile approach for targeted delivery of viral vectors that should assist in the study of cell types and circuits in the cerebral cortex.

## Introduction

The rapid expansion of genetic and optical tools for monitoring and manipulating cells in the rodent brain has redefined how neuroscientists study brain structure and function. For example, neuroscientists often employ adeno-associated viruses (AAVs) to genetically modify brain cells to make them amenable for imaging^1^. However the delivery of these vectors, which typically relies on blood brain barrier (BBB) permeable AAVs or direct micro-injection can be spatially imprecise, technically challenging with risk of infection, haemorrhage and worst of all, damaging to the same regions one intends to image.

There are several new tools available that enable minimally invasive expression of AAVs in the rodent brain. The recent development of blood brain barrier permeable AAVs with cell type specific promoters, provide a new alternative for widespread expression of specific proteins in the brain without the need for micro-injections^2,3^. For more focal and precise expression of proteins, Yao et al.,^4^ created light inducible recombinases that can be activated *in vivo* with single and two-photon light sources. While revolutionary, the application of these methods has been slowed by the fact that delivery of the light sensitive cre-recombinase involves the aforementioned BBB permeable AAVs, which can yield capricious expression in some mouse strains or avoid certain cells *in vivo*^5^. Another recent approach that has generated tremendous excitement is the use of focused ultrasound (FUS) to remotely and transiently disrupt the BBB with microbubbles, in order to deliver AAVs^6–9^. The benefits of this method is that one can non-invasively deliver AAVs to any brain region of interest. However, some limitations are the need for potentially expensive equipment to implement FUS, the inability to control the extent of transfection on a micrometer scale, and the unavoidable sterile inflammation found within the volume of tissue targeted by FUS. While all these different approaches have enormous potential for minimally invasive, spatially targeted delivery of AAVs or expression of cre-dependent proteins, they are not ideally suited or sufficiently simple for all experimental applications.

To address this need, we have optimized a simple, yet effective and titrate-able method to achieve targeted, ultra sparse AAV transfection of cortical neurons and astrocytes in the cerebral cortex. This facile approach involves the intravenous administration of commercially available AAVs followed by optically puncturing single capillaries with the same femtosecond laser used to image cells *in vivo*. Since the dose of AAVs can be titrated, the extent of cellular transfection (tdTomato reporter, GCaMP, ChR2) can be manipulated. Furthermore, the extent of inflammation and putative tissue damage is extremely limited compared to traditional micro-injection procedure, thereby allowing one to image cells at the target site with minimal optical distortion that invariably accompanies tissue damage (eg. Edema).

## Materials and Methods

### Animals

Two to 12 month old male mice on a C57BL/6J background were used in this study. For experiments involving conditional cre-dependent expression of TdTomato, we utilized Ai9 reporter mice (B6.Cg-*Gt(ROSA)26Sor*^*tm9(CAG-tdTomato)Hze*^/J, JAX# 007909) crossed with a microglia specific cre driver line (*Tmem119*^*em1(cre/ERT2)Gfng*^/J, JAX# 031820). In accordance with our ethical obligation to reduce animal numbers, some of these mice were re-used from another imaging study (unpublished study imaging microglia). We should note these mice were used as genetic controls in the previous study and had been given at least 8 weeks rest before being incorporated in the present study. Imaging of cre-dependent expression of eYFP tagged channelrhodopsin was achieved using Ai32 mice (B6.Cg-*Gt(ROSA)26Sor*^*tm32(CAG-COP4*H134R/EYFP)Hze*^/J; JAX# 024109). And finally, constitutive eGFP expressing microglia were imaged using heterozygous Cx3cr1-eGFP mice (JAX # 005582). All mice were group housed (when possible) on a 12 hour light/dark cycle in ventilated racks in a humidity (RH 40-55%) and temperature controlled room (21-23ºC). Mice were provided food and water *ad libitum*. All experiments comply with the guidelines set by the Canadian Council on Animal Care and approved by the local university Animal Care Committee. Reporting of this work complies with ARRIVE guidelines.

### Cranial window surgery

All mice imaged in the present study had a craniectomy based cranial window implanted over the right hemisphere. As previously described^20-21^, two to four month old mice were anesthetized with isoflurane (2% for induction and 1 – 1.3% for maintenance) mixed in medical air (flow rate: 0.7L/mL) and then fitted into a custom head fixing plate for surgery. Body temperature was maintained at 37°C with a feedback based heating system. After local injection of topical anesthetic, the scalp was cut and retracted. First, a 12 mm diameter titanium ring (used to hold the head during imaging, 7 mm inner diameter and 1.75 mm thick) was affixed to the skull with metabond adhesive. The skull was carefully thinned in a circular manner (∼4mm diameter) with a dental drill until the bone became transparent. Fine foreceps were used to lift the skull flap off and ice cold HEPES-buffered artificial cerebral spinal fluid (ACSF) was applied to the cortical surface. A 5 or 6mm circular coverslip was placed over the exposed brain and glued into place using cyanoacrylate glue and dental cement. Mice were allowed to recover on top of a heating blanket and then returned to their home cage.

### Intrinsic optical signal (IOS) imaging

For IOS imaging, mice were lightly anesthetized with isoflurane (1% in medical air) and mounted on an upright Olympus microscope with body temperature maintained at 37°C. First, high contrast images of the surface vasculature were generated by illuminating the surface with white light and collecting reflected light through a X2 objective (NA=0.14) and a YFP emission filter. For each IOS imaging trial, we focused a red LED light (635nm, ∼20mW at back aperture) 200-300µm below the cortical surface to minimize the contribution of large surface vessels. A total of 12-36 trials were collected per limb, with each stimulation trial followed by a no-stimulation trial. Each 3s trial consisted of 1s of baseline followed by 1s of stimulation (or no stimulation). A piezoelectric wafer (Piezo systems, Q220-A4-203YB) was used to stimulate the contralateral forepaw or hindpaw for 1 sec at 100Hz using 5ms bi-phasic pulses. Twelve-bit image frames (376×252 pixels, 16.7µm / pixel) collected all reflected red light at a frame rate of 100Hz using a MiCAM02 camera (Brain Vision). Using Image J software (version 1.53), the 12-36 imaging trials were averaged together and mean filtered (5 pixel radius). Stimulation induced changes in reflected light (IOS signal shown as ΔR/Ro) were calculated by subtracting and then dividing all images to an average intensity projection of pre-stimulus images (Ro). Maps of the forelimb or hindlimb somatosensory cortex were created by thresholding ΔR/Ro values at 70% of maximum intensity and superimposed on the cortical surface.

### *In vivo* two-photon imaging and targeted delivery of AAVs to cortical regions

All mice implanted with a cranial window were allowed to recover for at least 5 weeks before experiments commenced. Mice were anesthetized with isoflurane mixed with medical air (1%) and body temperature was maintained at 37°C. Their head was fixed in place with a custom head holding apparatus. Two-photon images were collected with a 40X water immersion IR objective (Olympus, NA=0.8) using an Olympus FV1000MPE laser scanning microscope coupled to a mode-locked Ti:sapphire femtosecond laser. For imaging Fluorescein (FITC), eGFP, eYFP ChR2, or tdtomato/GCaMP6s, the laser was tuned to the following wavelengths: 805, 900, 920 or 945nm, respectively. Simultaneous imaging of tdtomato and FITC dextran was achieved using 945 nm wavelength.

To visualize the cerebral vasculature and allow for subsequent puncture of targeted capillaries, we made up 0.1mL solution of 2.5-5% FITC-dextran (70kDa, Sigma-Aldrich #46945) dissolved in sterile saline solution. Various AAVs were then added to the FITC dextran solution and administered intravenously through tail vein or retro-orbital injection. The following AAVs were injected: 1) AAV1.hSyn.cre.WPRE.hGh (Addgene #105553) at one of three doses (1.73×10^12^, 6.92×10^12^ and 1.38×10^13^ GC/kg), 2) AAV1.CAG.tdTomato (5.06×10^12^ GC/kg; Addgene #59462), 3) AAV1.hSyn.GCamP6s.WPRE.SV40 (6.67×10^12^ GC/kg, Addgene #100843). Before puncturing a capillary of interest, we collected a baseline image stack of the cortical vasculature at 2μm Z-steps covering an area of 317 × 317μm (1024×1024 pixels, 0.31μm/pixel). A flowing capillary (∼3-7μm in width), 50–250μm from the pial surface was selected. Rupture was induced with our femtosecond laser, and monitored in real time with focal high power raster scanning over a region of interest (805nm, ∼390mW at back aperture, 4-5μm diameter region centered on the vessel) for 3-6 six seconds. In order to maximize sampling, we typically ruptured 4-8 capillaries per mouse, spaced at least 500 µm apart from each other. To confirm AAV mediated expression of biosensors or reporter proteins, we re-imaged a volume of cortical vasculature and AAV labelled cells (∼0.01-0.03mm^3^, see image stack parameters above) 2-5 weeks after capillary rupture. For GCamP6s imaging, single plane images were collected at 4 or 8Hz covering an area of 158.4×158.4µm. Sensory evoked calcium transients were elicited with vibrotactile stimulation of the contralateral limb at 100Hz for 1.5s, starting 5s into each 10s trial, and repeated over 6-10 trials. Sensory evoked GCaMP6s signals were calculated by extracting fluorescence values from the soma of interest. Neuropil fluorescence surrounding each cell was subtracted from the soma fluorescence to estimate “true” soma fluorescence. Corrected soma GCaMP6s fluorescence was then subtracted and divided by pre-stimulus soma fluorescence (Fo was defined as median F value before stimulus) to yield a percent ΔF/Fo value.

### Statistical analysis

Statistical analysis of the data was conducted in Microsoft Office Excel or GraphPad Prism. Data sets were first checked for normality and outliers were identified using a ROUT test set at 1%. The non-parametric Kruskal-Wallis statistic with post-hoc Dunn’s or Mann-Whitney test were used to analyze dose dependent differences in cell labelling. Cell type specific differences in the distance from rupture site were assessed with an unpaired t–test. Linear regression was used to test the relationship between the extent of cell labelling and diameter of punctured capillaries. All p values ≤ 0.05 were considered statistically significant. All the data are presented as mean ± standard error.

## Results

Conventional delivery of AAVs using micro-injection is technically challenging and inevitably leads to considerable tissue damage associated with the micro-pippettes. As an alternative, we considered the possibility that intravenous injection of an AAV followed by laser based perforation of a capillary could provide precise and minimally invasive delivery. Our rationale was based in part, on the fact that sparse cre-recombinase dependent reporter expression in the brain can be achieved with direct micro-injection of a very dilute solution of virus (eg. 1:20,000 dilution, see **Supp. Fig. 1**). Given the blood volume of an adult mouse is approximately 1.5-2.5mL, it was reasonable to think that a comparable dilution could be attained with an intravenous injection of a high titre AAV. Therefore we intravenously injected AAV1.hSyn.cre.WPRE.hGh (Addgene #105553, 6.92×10^12^ GC/kg) diluted in 2.5-5% FITC dextran (70kDa; Sigma-Aldrich #46945) into adult mice implanted with a cranial window (**Fig. 1A**), that conditionally expresses the fluorescent reporter tdtomato (Ai9, JAX# 007909). To precisely target cells within a specific cortical region, we optically perforated a single capillary between 50 and 250µm below the cortical surface with our high power femtosecond laser. Puncture of a capillary was easily confirmed by the appearance of extravascular dye fluorescence surrounding the rupture (**Fig. 1B**). Re-imaging the same region 2 and 4 weeks later revealed sparsely labelled neurons and astrocytes adjacent to the ruptured capillary (**Fig. 1C and D**). Our success rate in achieving cre-dependent tdtomato expression was 95.6% (22/23 ruptures in 4 mice). We should note that since we ruptured capillaries in multiple cortical regions (≥500µm from each other) over the span of 60 min after AAV injection, we did not find any time dependent decrement in successful AAV mediated cell labelling. To prove this approach could be applied to other cre-dependent strains, we injected AAV1.hSyn.cre (6.93×10^12^ GC/kg) into mice that conditionally express YFP tagged channelrhodopsin-2 (ChR2; Ai32, JAX #024109). Doing so led to ChR2 expression in neurons and astrocytes next to the ruptured capillary (**Fig. 1E-F**; 100% success rate from 11 ruptures in 2 mice; Ai32, JAX #024109).

**Figure 1.**
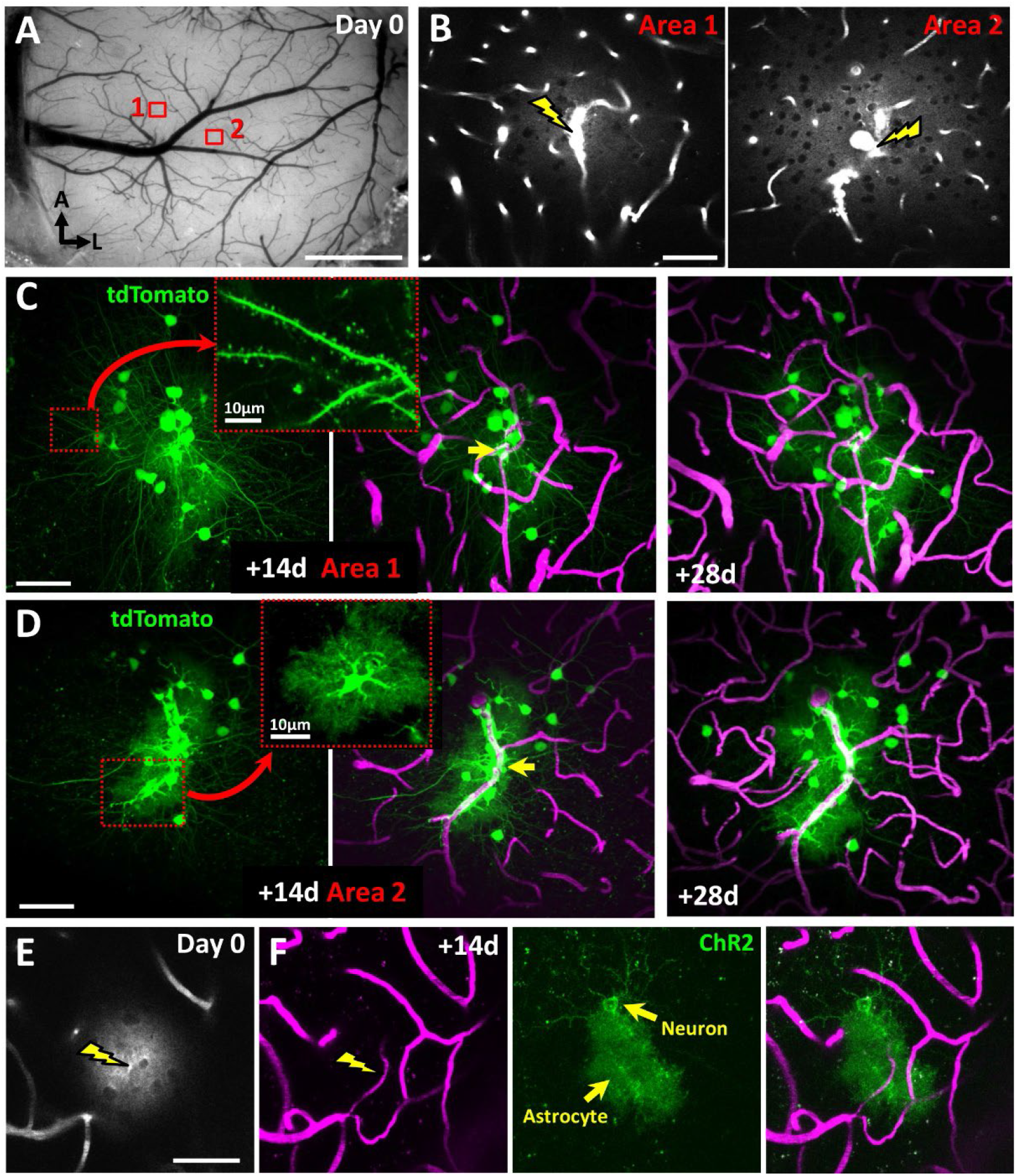
Targeted delivery of AAVs and sparse expression of cre-recombinase dependent fluorescent proteins in cortical neurons. **A:** Brightfield image of the cortical surface in an Ai9 reporter mouse implanted with a cranial window and intravenously injected with AAV1.hSyn.cre. Note the 2 areas where a single capillary was ruptured in each (denoted by lightning bolt). **B:** Two-photon images showing extravasation of fluorescently labelled blood plasma in Areas 1 and 2 immediately after rupturing a capillary. **C and D:** *In vivo* maximal intensity z-projection images taken 14 and 28 days (left and right panels, respectively) after vessel rupture in Areas 1 and 2, showing the cre-dependent expression of tdtomato in nearby cells (green) and vasculature labelled with fluorescent dye (magenta). Note the bright tdTomato signal that allows visualization of fine dendritic structure or peri-vascular astrocytes (see insets in C and D, respectively). **E:** Image showing extravasation of plasma dye immediately after vessel rupture. **F:** *In vivo* maximal intensity z-projection images taken 14 days after vessel rupture revealing the Cre-dependent expression of eYFP labelled ChR2(H134R) in an Ai32 mouse. Scale bars in A: 1mm; B-E: 50µm, insets in C and D: 10µm.

Next we wanted to determine if other AAVs (such as constitutive ones), could be delivered and express their payload, without the need for a cre-dependent mouse strain. Intravenous injection of AAV1.CAG.tdTomato (5.06×10^12^ GC/kg; Addgene #59462) followed by capillary perforation induced tdtomato expression in neurons and astrocytes 2-3 weeks later (**Fig. 2A**; 100% success in 15 ruptures from 2 mice). Although we could detect labelled cells in each experiment, the brightness of tdTomato expression was considerably lower than the Cre-dependent expression of tdTomato in the Ai9 reporter strain (using the same excitation wavelength and laser power). In our next set of experiments, we tested a comparable dose of AAV1.hSyn.GCamP6s.WPRE.SV40 (6.67×10^12^ GC/kg, Addgene #100843). In this case, we functionally mapped the forelimb and hindlimb somatosensory cortex using intrinsic signal optical imaging and targeted single capillaries in these regions (**Fig. 2B,C**). Two to three weeks later, we could detect GCamP6s expressing neurons and astrocytes near the site of rupture (88% success from 25 ruptures in 4 mice; **Fig. 2C**). To determine if these cells were viable and active, we longitudinally imaged neuronal calcium transients in response to 1s of vibrotactile stimulation of the contralateral limb (**Fig. 2D**). As shown in **Figure 2E**, tactile stimulation reliably evoked calcium transients over multiple weeks. Importantly the long term sensory responsiveness of these neurons indicate that the cells in the immediate vicinity of the perforated capillary remain functional and appear to suffer no ill effects from the transient rupture. Collectively, these experiments indicate that constitutive AAVs (ie. non-Cre recombinase dependent) can be delivered and expressed in the mouse cortex using our approach.

**Figure 2.**
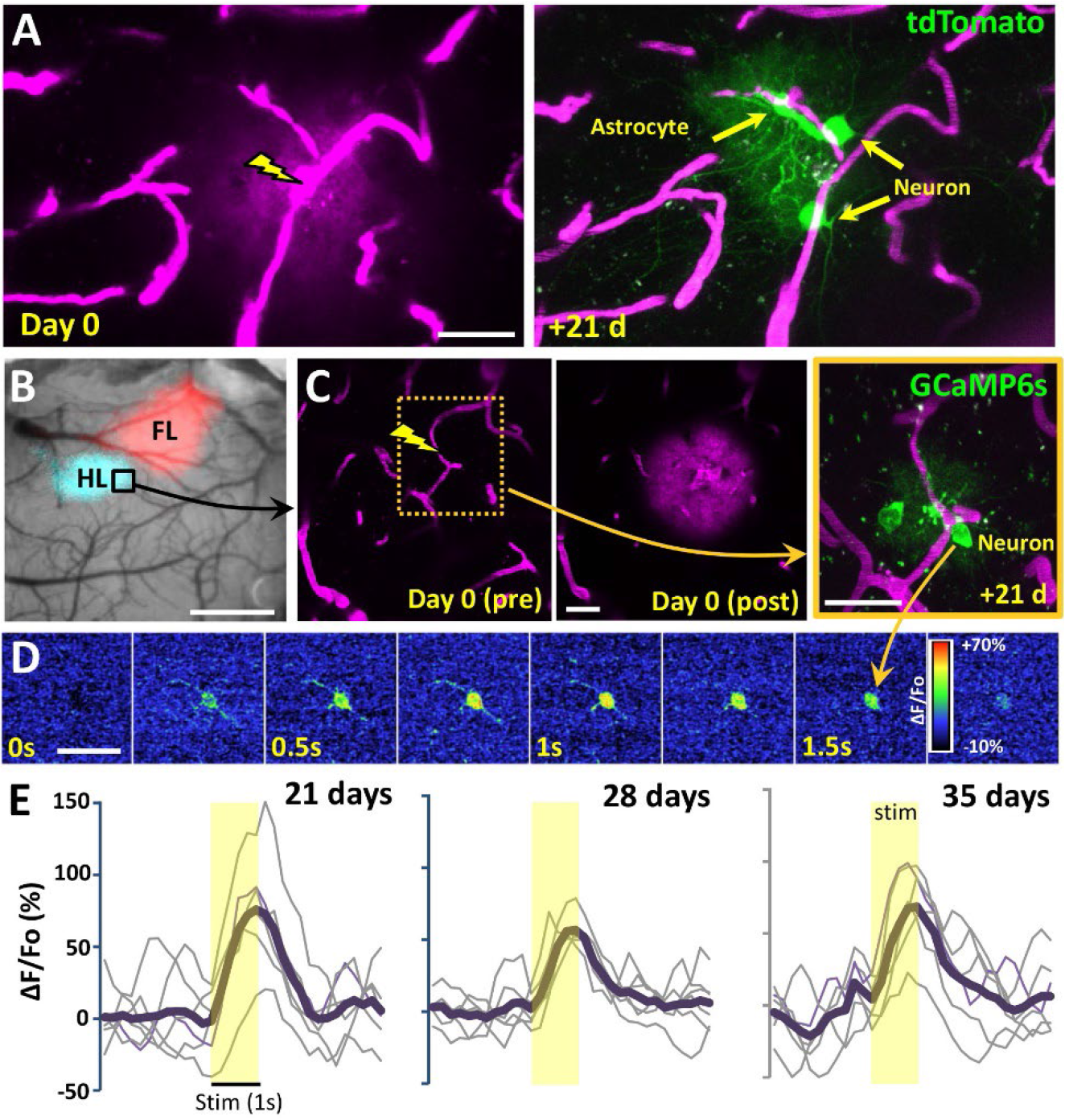
Targeted delivery and expression of constitutive (non-cre dependent) AAVs to cortical neurons. **A:** Two-photon maximum intensity projection images showing rupture of a cortical capillary after intravenous injection of constitutive AAV (AAV1.CAG.tdTomato) and resultant expression of tdTomato in cortical astrocytes and neurons 2 weeks later. **B:** Brightfield image showing the cortical surface with intrinsic optical signal derived maps of the forelimb (FL) and hindlimb (HL) primary somatosensory cortex superimposed on top. **C:** Two-photon images show a capillary immediately before and after rupture on day 0, as well as resulting GCaMP6s expression in nearby cells 21 days later. **D:** Color montage illustrates time course of hindlimb evoked neuronal calcium responses (average of 6 stimulation trials, expressed as % ΔF/Fo). **E:** Individual and averaged (6 trials, thick black line) calcium responses (from neuron shown in C and D) following 1s vibrotactile stimulation of the contralateral hindlimb, collected at 21, 28 and 35 days after vessel rupture. Scale bars in A,C,D: 30µm; B: 1mm.

For imaging and understanding the wiring diagram of cortical neurons at different scales, it would be helpful to titer AAV mediated expression in targeted regions. Therefore we intravenously injected 3 different doses of AAV1.hSyn.cre.WPRE.hGh in Ai9 tdtomato reporter mice. Comparison of reporter expression 2 weeks after capillary perforation indicated a dose dependent increase in cells expressing the tdtomato reporter (**Fig. 3A,B**; Kruskal-Wallis test, Main effect of dose: p<0.0001). At the lowest dose (1X: 1.73×10^12^ GC/kg), tdtomato labelled cells were found at 24 of 26 sites (92.3% success) with a median of 2 cells per site (**Fig. 3B**). With higher doses, the success rate increased (95.6 and 100% for 4X and 8X, respectively) as did the median number of labelled cells per site (left panel in **Fig. 3B**; 6 and 17 cells/site for 4 and 8X dose, respectively). By examining the morphology of labelled cells (right panel in **Fig. 3B**), the proportion of neuronal vs astroglial cells labelled was not significantly different at 1 and 4X doses, although the highest dose labelled significantly more neurons than astrocytes. Next, we examined the proximity of labelled cells to the rupture site. Our analysis shows that on average, neurons were located 60.62µm away, whereas astrocytes were significantly closer at an average distance of 38.93µm (**Fig. 3C**). And finally, since capillaries can vary in diameter (from 3-8µm), we plotted the number of labelled cells per site as a function of lumen diameter (**Fig. 3D**). Linear regression analysis indicated a significant relationship (R^2^=0.137, p=0.014) suggesting that rupturing larger capillaries tends to label more AAV infected cells. In summary, these results show that AAV mediated transfection of cortical cells leads to spatially localized expression that can be tittered by dose and the size of the vessel perforated.

**Figure 3.**
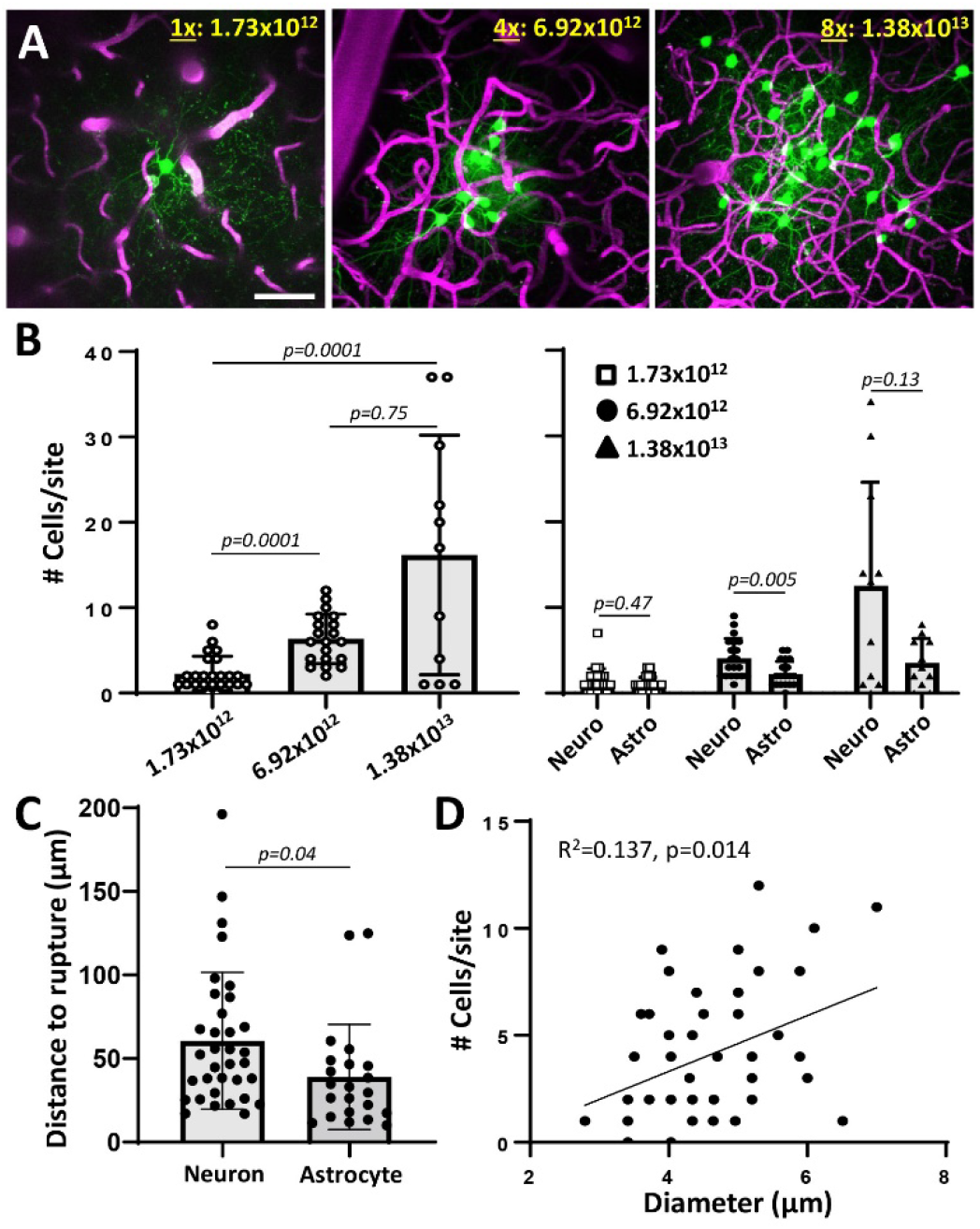
Tittered expression of cre-dependent AAVs in cortical neurons allow for ultra-sparse labelling applications. **A:** *In vivo* maximum intensity z**-**projection images collected from an Ai9 tdTomato reporter mice 2 weeks after intravenous injection of AAV1.hSyn.Cre and targeted rupture of a capillary. Note the increase in TdTomato expressing cells from low to higher doses of AAV (1-8X, AAV dose expressed GC/kg). **B**, Left panel: Bar graphs show a significant effect of AAV dose on the number of cells labelled per site (Kruskal-Wallis statistic: 20.11, p<0.0001). Right panel: the number of neurons or astrocytes per site as a function of AAV dose. **C:** graph shows that neurons were located slightly, but significantly further form the site of rupture than astrocytes. **D:** linear regression showing relationship between the diameter of ruptured capillaries and the number of labelled cells. Statistics based on Kruskal-Wallis statistic with post-hoc Dunn’s (B: left panel) or Mann-Whitney (B: right); unpaired t-test (C); linear regression (D). Data are presented as mean ± standard error. Scale bar in A: 50µm.

While there are many possible applications for this method, we highlight one example focused on cortical plasticity following stroke. Our lab and several others have used longitudinal 2-photon imaging through a cranial window to describe structural and functional changes to cortical circuits in the days and weeks that follow an ischemic stroke in the forelimb somatosensory cortex^20-21^. An obvious, yet until now very difficult experiment, would be to functionally identify the part of the forelimb cortex that emerges weeks after stroke (so called “re-organized” or “re-emergent” cortical representation), and use AAVs to image and/or map their connections. Using a conventional micro-injection approach would be problematic because it lacks precision and would cause further damage to peri-infarct tissues, which are already vulnerable to insults, especially the vasculature. As shown in Figure 4, we identified the forelimb primary somatosensory cortex before and after photothrombotic stroke using intrinsic optical signal imaging (**Fig. 4A**), and then targeted AAV mediated tdTomato expression to peri-infarct cells (**Fig. 4B,C**). Importantly, we did not see overt signs of tissue damage in the form of generalized vessel loss or abnormal permeability of plasma dye across the BBB in subsequent imaging sessions (**Fig. 4D**). The fact that the majority of ruptured capillaries recover and recanalize after 2 weeks (see examples in **Fig. 1E**, 2**A**,**B** and **4D**) agrees with previous work from our lab^14^, and also correlates with the resolution of microglia related inflammation around the rupture site (**Supp. Fig. 2**).

**Figure 4.**
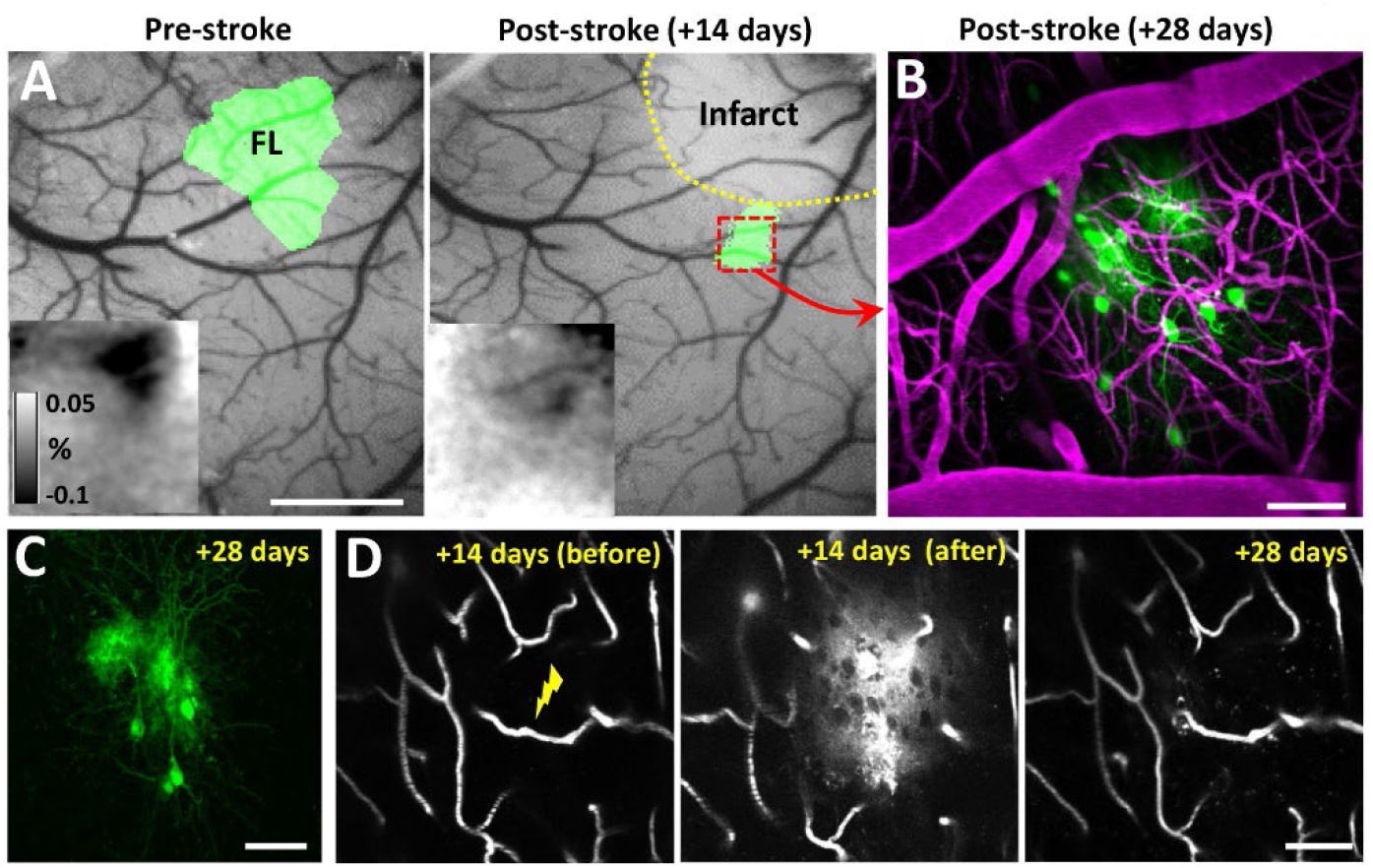
Application of the method for examining structural plasticity in cortical neurons within functionally altered somatosensory maps after stroke. **A:** Brightfield images of the cortical surface overlaid with intrinsic optical signal maps of the forelimb somatosensory cortex, before and 14 days after focal ischemic stroke. Insets show intrinsic optical signal reflectance maps (% dR/Ro) generated from stimulating the contralateral forelimb. **B:** *In vivo* maximum intensity projection image showing tdtomato expressing cells within the remaining portion of forelimb map, 2 weeks after induction of vessel rupture and intravenous injection of AAV1.hSyn.Cre (6.92×10^12^ GC/kg; see middle panel in D). **C:** Post-mortem confocal image (from coronal brain section) showing tdtomato labelled cells displayed in B. **D:** Two-photon images showing the targeted capillary immediately before (left) or after (middle) rupture. Imaging the same capillary 2 weeks later (right, +28 days) shows it was preserved and regained blood flow. Scale bars in A: 1mm; B-D: 50µm.

## Discussion

Here we have validated a simple, minimally invasive approach for near single cell targeted AAV expression in the mouse cortex. This method leverages a common tool in neuroscience laboratories, the 2 photon microscope, which most labs interested in optical reporters and actuators already have and use. We exploited the fundamental advantage of multi-photon excitation which is spatially restricted to a focal point of excitation^10,11^, to perform targeted perforations of cortical capillaries. When combined with AAVs injected into the bloodstream, this transient rupture allows an extremely small quantity of viral particles into the cortical parenchyma and transfection of only a few adjacent neurons and astrocytes (Figure 1 and 3). We further demonstrate this method is an effective tool for targeted and limited expression of different AAVs and genetic payloads (Figure 2). We also have shown that by varying the dose of AAV and the diameter of vessels targeted, one can titrate the level of expression from just one or two cells to several dozen. Lastly, we demonstrate a practical application of this methodology, driving AAV dependent expression of tdTomato in sparse set of surviving peri-infarct neurons (Figure 4). The advantages of this methodology are its ability to precisely deliver AAVs with micron-level precision and sparsely label neurons at a density of one’s choosing without risking damage associated with direct micro-injection.

Direct micro-injections remain the most common method for gene transduction in the brain using viral vectors, especially when the goal is focal uptake of an AAV and expression of the transgene. This method is effective, relatively simple and cost effective but remains limited for selective targeting. Titrating viral loads and thus expression, with glass pipettes requires either excessive dilutions or extremely small volumes. Neither of these strategies circumvent the inherent lack of precision of inserting a ∼5-200μm wide glass pipette (from tip to further up the bevel) into the cortex. Additionally the insertion inevitably leaves a path of damage and AAV backflow up the insertion track and less precise expression (Supplemental Fig 1.). The method described in the present study is not the only approach to improve upon the traditional method of AAV delivery by insertion of a glass pipette. For example, recent studies have proven that focused ultrasound combined with intravenously injected microbubbles, can deliver systemically administered AAVs to any brain region of interest^7,12^. While this approach could be a transformative step for clinical application of gene therapy, BBB disruption and gene transduction occur over a relatively large volume (∼0.125-1 mm^3^), and therefore it is not suitable for micron level precision of AAV delivery and longitudinal imaging of sparsely labelled cells. Another major innovation was the development of light inducible Cre recombinases^4^. These constructs work along similar principles, the transgene of interest is widely expressed, usually by injection of blood brain barrier permeable AAV-PHP^13^, followed by focal application of light to activate Cre recombinase. A significant advantage of this method is that it could, in theory, allow for ultra-sparse and circuit specific manipulations by chaining different intersectional criteria, such as cell specific Cre expression, cell specific transgene and the spatial / temporal application of light. While this method holds tremendous potential, the ability to target cells within a very specific region of interest is dependent, and conceivably limited by how well the existing AAV-PHPs can deliver the light inducible recombinase. For example, it is known that efficiency of AAV-PHP transduction is variable and dependent on factors such as mouse strain^2,5^ and exhibits a tropism for certain neurons (eg. Pyramidal neurons in cortical layers 2/3 and 5), although cell specificity has been improving. To obviate this issue, direct micro-injection of AAV-PHP has been used^4^, although for reasons previously stated, the invasive effects of direct injection are less than ideal for *in vivo* imaging.

While we believe the approach described in the present paper will be useful to the multiphoton imaging community, there are important limitations that should be considered. Firstly, our approach is not completely non-invasive since we still have to optically puncture a microvessel for AAV delivery. However, we show that cells in the immediate vicinity of the ruptured capillary display preserved activity patterns, namely sensory evoked responsiveness over weeks time (see **Fig 2B-E**). Furthermore, punctured vessels usually recover completely (**Fig. 1E**, 2**A**,**B** and **4D**) and highly local inflammatory microglial responses subside within 2 weeks. A second limitation is we only tested AAV serotype 1, and in mice with a C57 background. Thus we cannot guarantee that this approach will work for all AAVs or animals tested. However we do note that our capillary perforation delivery of AAV.syn.cre worked well in both Ai9 and Ai32 reporter strains, suggesting this is a robust delivery method. Furthermore, the delivery of these viruses is likely based on passive diffusion through the ruptured vessel for at least 30 minutes after rupture (but not more than 24 hours)^14,15^, rather than an active receptor based transport (eg. LY6A receptor needed for AAV-PHP) which can limit AAV delivery in certain mouse strains. Another limitation is that systemic administration of AAV and uptake of virus throughout the body, raises the possibility of organ toxicity. Although our mice did not display signs of morbidity with any AAV dose tested (pain or sickness related behaviours) for at least 4-6 weeks after injection, future refinements could incorporate intranasal delivery of AAVs which leads to viral expression in the brain but with significantly lower bio-distribution in peripheral organs^12^. We should also note that we did attempt our method in 4 mice that were previously injected with an AAV (> 6 weeks prior). However, these attempts were unsuccessful in all 4 mice likely due to the production of AAV neutralising antibodies, which has been reported in other studies with different AAVs^16^. And finally, the present method is limited by the need for a cranial window and inherent depth limitations of 2-photon imaging, particularly given the high laser powers required for vessel perforation^17,18^. While light scattering in tissue is a fundamental limit of any optical method, these concerns can be managed by surgical expertise, choice of cranial window (open vs thinned skull) and emerging deep tissue imaging methods such as 3 photon imaging and GRIN lens^19^. Presumably, any laboratory that is currently using 2-photon microscopy for *in vivo* imaging could easily apply this technique to their study with minimal cost and no need for additional equipment.

Any new approach is only as useful as it’s best applications. Here we used this approach to address a challenge we have considered for years; how to non-invasively label a sparse set of surviving peri-infarct neurons within small fragments of the stroke affected somatosensory cortex. While we have simply shown proof of concept here, our approach allows us to target surviving neurons, whose location is notoriously difficult to predict after stroke, and unambiguously trace these neuronal projections across the brain. Moreover, we can restrict AAV delivered opto-or chemo-genetic actuators to neurons in these surviving regions, so that we can precisely manipulate their function/activity and potential role in stroke recovery, with simple, imprecise light sources (eg. Surface LEDs) or systemic administration of chemogenetic ligands^20,21^. The spatial resolution of this method is also advantageous for investigating finely organised topological maps, such as targeting functional subdomains within a single whisker barrel^22^, or retinotopic/feature-specific microdomains in the visual cortex^23,24^. Several recent groundbreaking studies have used retrograde viral constructs to trace presynaptic inputs to a single post-synaptic neuron, infected by single cell delivery with a glass pipette^24,25^. This method is extremely challenging and potentially damages incident axons to the target neuron as the pipette is moved into a juxtasomal position. Therefore an optical approach for delivery of viral constructs to single neurons, could significantly improve the success rate of these experiments and lower the expertise needed to implement them.

## Acknowledgements

We are grateful to Angie Hentze and Taimei Yang for managing the mouse colony. We thank Stephanie Taylor and Eslam Mehina for microglia images. Work was supported by operating, salary and equipment grants to C.E.B. from the Canadian Institutes of Health Research (CIHR), Heart and Stroke Foundation (HSF), Natural Sciences and Engineering Research Council (NSERC).

## Supplementary Information to

**Supplementary Figure 1.**
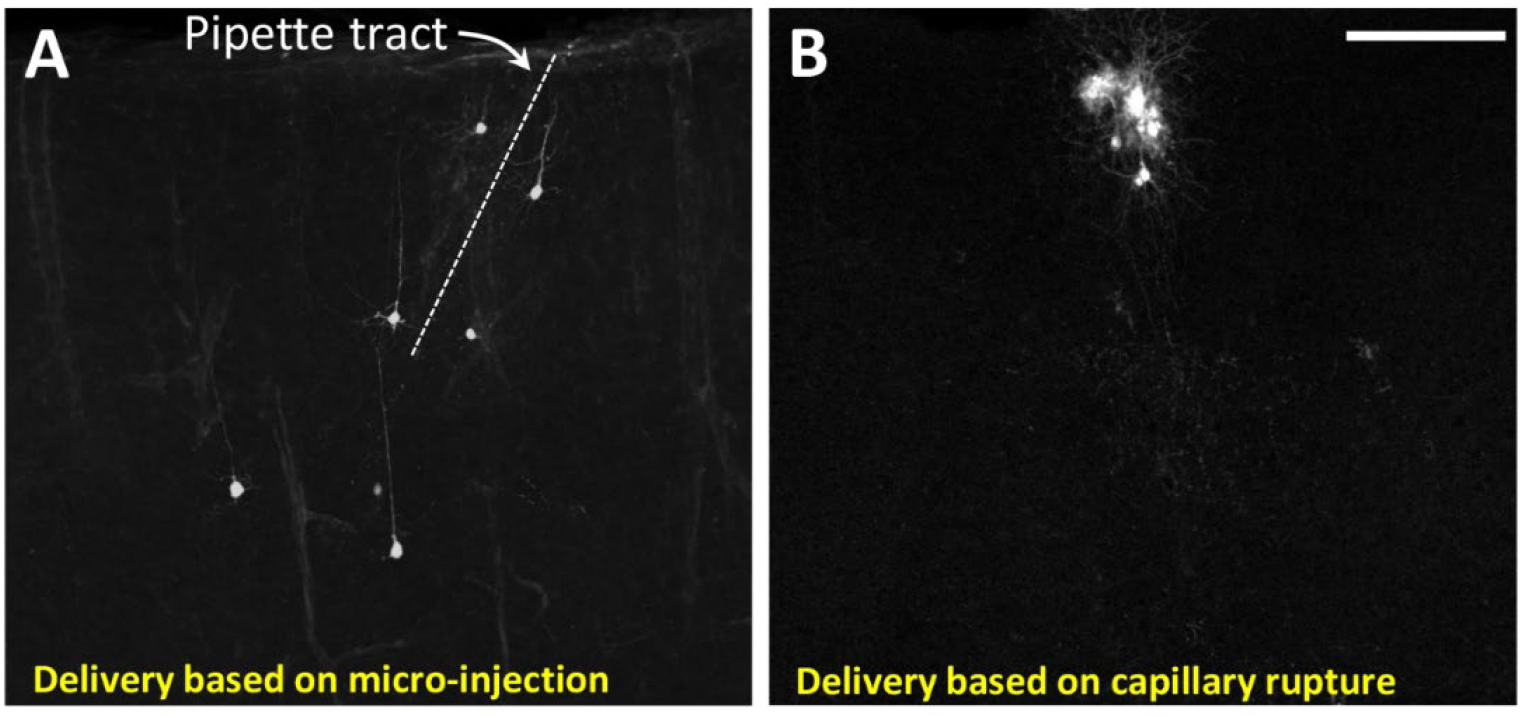
Comparing the spatial extent of cre-mediated labelling of cells following intracortical micro-injection of AAV.hSyn.cre versus rupture of a single capillary. Confocal images from a coronal brain section show conditional expression of tdtomato 3 weeks after AAV injection with a glass micropipette (A; 0.4uL, ∼1.3×10^10^ GCs) or after rupture of a capillary (B; i.v. injection of 6.92×10^12^ GCs/kg). Scale bar = 200µm.

**Supplementary Figure 2.**
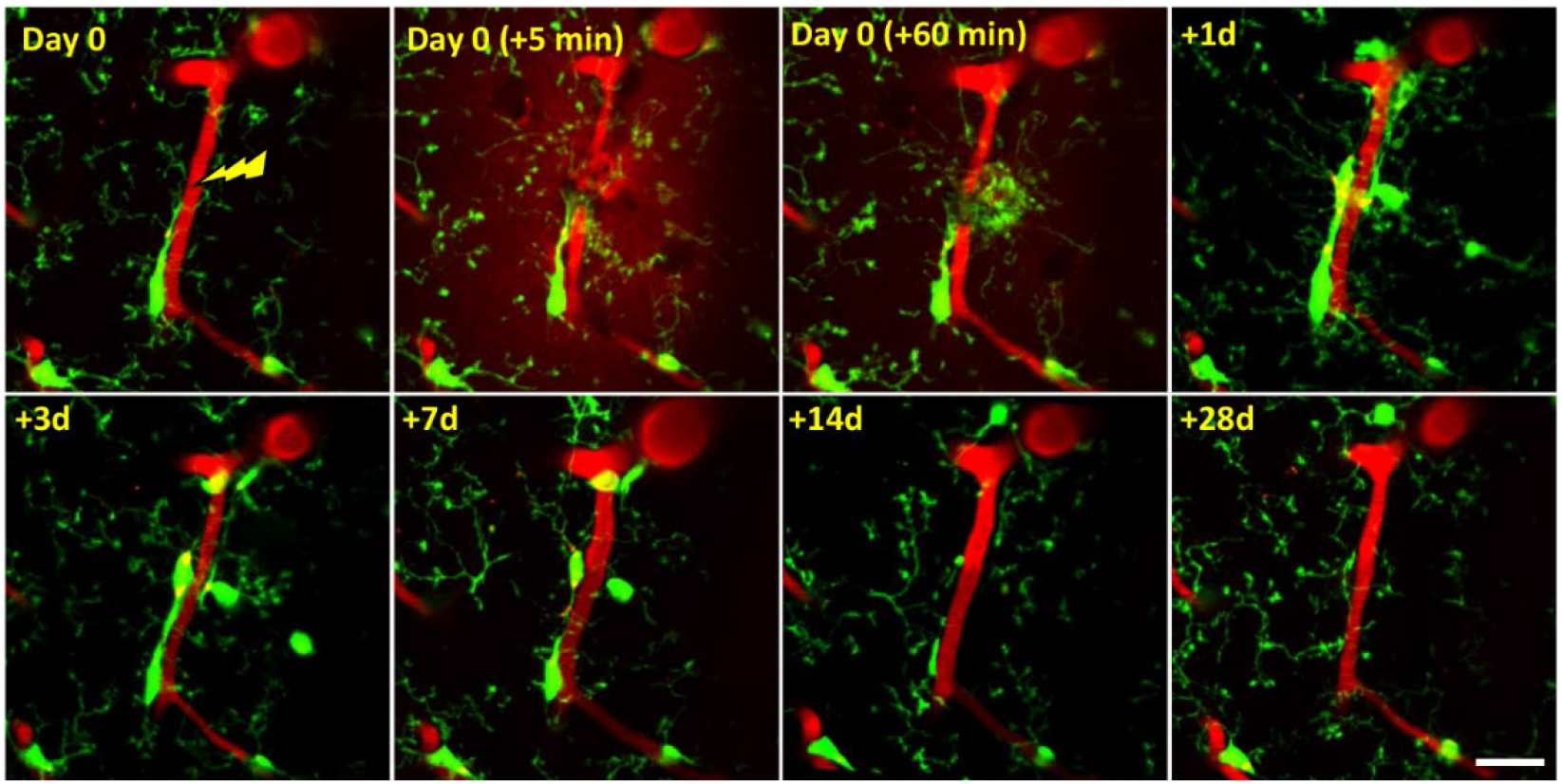
Time-dependent changes in microglial responses to capillary rupture. Longitudinal *in vivo* imaging of microglia in Cx3cr1^gfp/wt^ mice before and up to 28 days after rupture of a single capillary. Note the rapid accumulation of microglia processes around the bleed site and delayed recruitment of cells 1 day after injury. The capillary is retained and regains blood flow while microglial reactivity subsides over 14 days after rupture. Scale bar = 20µm.

## Notes

**Conflict of Interest Statement:** The authors have declared that no conflict of interest exists

### Competing Interest Statement

The authors have declared no competing interest.

