## Supplemental Data 1 for "Optical opening of the blood brain barrier for targeted and ultra-sparse viral infection of cells in the mouse cortex"

**Supplementary Figure 1. Comparing the spatial extent of cre-mediated labelling of cells following intracortical micro-injection of AAV.hSyn.cre versus rupture of a single capillary.** Confocal images from a coronal brain section show conditional expression of tdtomato 3 weeks after AAV injection with a glass micropipette (A; 0.4uL, ~$1.3 \times 10^{10}$ GCs) or after rupture of a capillary (B; i.v. injection of $6.92 \times 10^{12}$ GCs/kg). Scale bar = 200µm.

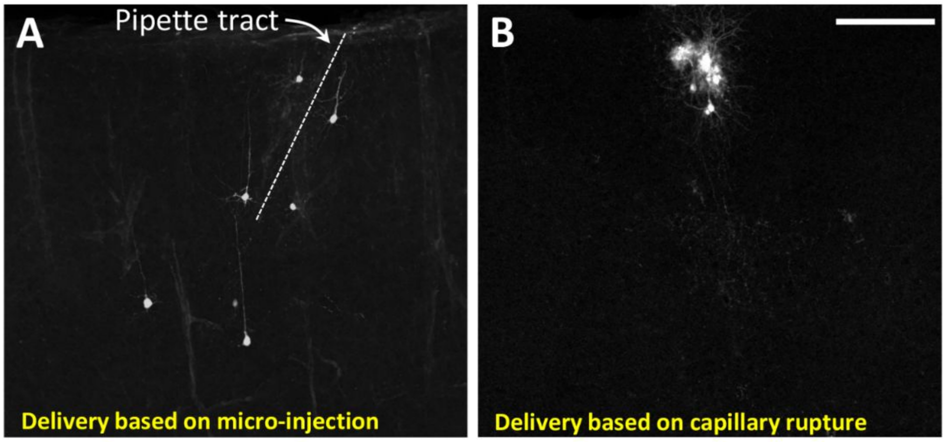


**Supplementary Figure 2. Time-dependent changes in microglial responses to capillary rupture.** Longitudinal *in vivo* imaging of microglia in $Cx3cr1^{gfp/wt}$ mice before and up to 28 days after rupture of a single capillary. Note the rapid accumulation of microglia processes around the bleed site and delayed recruitment of cells 1 day after injury. The capillary is retained and regains blood flow while microglial reactivity subsides over 14 days after rupture. Scale bar = 20µm.

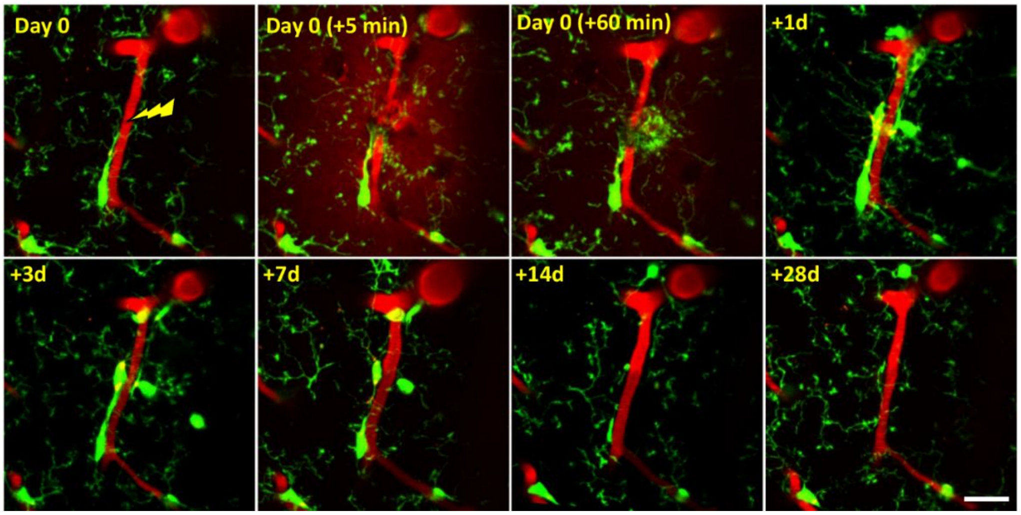
